# CA2 neurons show abnormal responses to social stimuli in a rat model of Fragile X syndrome

**DOI:** 10.1101/2025.09.10.674743

**Authors:** Margaret M. Donahue, Emma Robson, Alyssa M. Marron, Edel J. Fernandez, Misty Hill, Alexandra J. Mably, John B. Trimper, Darrin H. Brager, Laura Lee Colgin

## Abstract

Fragile X Syndrome (FXS) is a neurodevelopmental disorder that is highly comorbid with autism spectrum disorders and can cause abnormal social behaviors. The CA2 subregion of the hippocampus is essential for social memory processing and social recognition. A social interaction induces changes in CA2 neuronal firing; however, it is unknown whether these changes are impaired in FXS models. Here, we examined CA2 activity in a rat model of Fragile X Syndrome (*Fmr1* knockout rats). In *Fmr1* knockout rats, we observed impaired CA2 cell responses to social stimuli, despite similar social behaviors. Further, in CA2 of *Fmr1* knockout rats, we found reduced expression of oxytocin receptors and impaired whole cell responses to oxytocin. Together, these results raise the possibility that abnormal CA2 activity contributes to impaired social behavior in FXS and may suggest novel treatment targets for FXS patients.

**Significance statement:** Fragile X Syndrome (FXS) is a neurodevelopmental disorder that can result in abnormal social behaviors, including social avoidance. Activity in the CA2 subregion of the hippocampus is believed to support social recognition and social cognition. Yet, the extent to which the CA2 subregion of the hippocampus is affected by FXS is poorly understood. In this study, we identified specific impairments in CA2 neuronal responses to social stimuli in a rat model of FXS. Further, we provide evidence suggesting that CA2 responses to oxytocin, a neuropeptide released during social interactions, are abnormal in FXS.

## Introduction

Fragile X Syndrome (FXS) is a neurodevelopmental disorder that is highly co-morbid with autism spectrum disorders. Patients with FXS often display abnormal social behaviors, including social avoidance (Kau et al., 2004; Kaufmann et al., 2004; Lesniak-Karpiak et al., 2003; Merenstein et al., 1996; Williams et al., 2014). FXS is caused by a deficit in the function of Fragile X Messenger Ribonucleoprotein (FMRP). FMRP is the product of the *Fmr1* gene, which is located on the X chromosome. FXS is caused by hypermethylation and transcriptional silencing of the *Fmr1* gene (Pieretti et al., 1991; Verkerk et al., 1991), leading to a complete lack of FMRP production. FMRP is an RNA-binding protein (Ashley et al., 1993; Siomi et al., 1993) believed to regulate translation and protein synthesis in neurons (Jin & Warren, 2000; Richter & Zhao, 2021). Mouse and rat models of FXS (hereafter termed “FXS mice” and “FXS rats”, respectively) have been developed by knocking out the homologous *fmr1* gene in these species (Bakker et al., 1994; Hamilton et al., 2014; Mientjes et al., 2006; Tian et al., 2017).

Social behavior deficits have been observed in FXS rodent models. FXS mice display more anxiety-like behaviors during social interactions (McNaughton et al., 2008; Spencer et al., 2005) and lower levels of social interaction (Liu & Smith, 2009; McNaughton et al., 2008; Mineur et al., 2006; Oddi et al., 2014) than wild-type (WT) mice. In addition, FXS mice and rats may be deficient in distinguishing between positive and negative (McNaughton et al., 2008), or familiar and novel (Tian et al., 2017), social interactions.

The CA2 region of the hippocampus is essential for social recognition (Oliva, 2022). Inactivating CA2 pyramidal cells impairs social memory and recognition (Hitti & Siegelbaum, 2014; Meira et al., 2018; Oliva et al., 2020; Stevenson & Caldwell, 2014). CA2, like all subregions of the rodent hippocampus, contains place cells, neurons that have spatial receptive fields called place fields (O’Keefe & Dostrovsky 1971). Place cells often alter their firing patterns in response to changes in sensory or cognitive input in a process referred to as “remapping” (Colgin et al., 2008). During “global” remapping, place fields may appear, disappear, or change location. During “rate” remapping, place fields do not move, but cells’ in-field firing rates change. Previous research has shown that the introduction of a social stimulus to a familiar environment induces global remapping in CA2 place cells (Alexander et al., 2016; Robson et al., 2025). Furthermore, this process appears to be driven by the olfactory content of social experiences (Robson et al., 2025). CA2 place cell responses to social stimuli may reflect a cellular mechanism underlying CA2’s contributions to social memory.

In this study, we recorded CA2 place cells in FXS rats and WT control rats freely exploring an open field arena. We analyzed changes in CA2 place cell firing patterns when stimuli with a social olfactory component were added to the arena. We found that the presentation of social odors was associated with global remapping in CA2 place cells in WT rats but not FXS rats. Receptors for oxytocin, a neuropeptide that regulates social behaviors and social memory (Froemke & Young 2021), are prominently expressed in CA2 (Mitre et al. 2016), and administration of an oxytocin receptor agonist potentiated excitatory postsynaptic currents (EPSCs) in CA2 neurons (Pagani et al. 2015). Thus, we hypothesized that abnormal CA2 neuronal responses to oxytocin contribute to deficient CA2 place cell remapping to social stimuli in FXS rats. In accord with this hypothesis, we observed reduced oxytocin receptor expression and impaired depolarizing responses to oxytocin in CA2 neurons in FXS rats. Together, these results suggest that CA2 cellular responses involved in social information processing may be impaired in FXS. The results also point to potential mechanisms that may contribute to abnormal social behaviors in FXS.

## Materials & Methods

### Subjects

The FXS rats used in this study were *fmr1*-knockout rats on a Sprague-Dawley background (SD-*Fmr1*-/null^tm1Sage^). Control rats used in this study were WT Sprague-Dawley rats. Both WT and FXS rats were obtained from Inotiv. As FXS is an X chromosome-linked disorder, the prevalence of FXS is higher in males, and symptoms tend to be more severe in male patients than female patients. Therefore, we chose to use male rats. All experiments were conducted according to the guidelines of the United States National Institutes of Health Guide for the Care and Use of Laboratory Animals. *In vivo* and *in vitro* studies were conducted under protocols approved by the Institutional Animal Care and Use Committees at the University of Texas at Austin and the University of Nevada at Las Vegas, respectively.

*Subjects for CA2 place cell recordings and open field exploration.* CA2 place cell recordings in this study were collected from twelve male rats. Six were WT control rats, and six were FXS rats. The six WT rats used in the *in vivo* portion of this study were included in one previously published study (Robson et al., 2025); one of the WT rats and three of the FXS rats were included in another previously published study (Donahue, Robson, & Colgin 2025). Rats were between the ages of 3 and 10 months old at the time of surgery. Before surgery, rats were double or tripled housed with littermates in same-genotype cages. Following surgery, rats were housed singly in custom-built acrylic cages (40 cm x 40 cm x 40 cm) to prevent damage to implants. Enrichment material (i.e., paper towel rolls, wooden blocks, etc.) was added to the cages, and rats were housed directly next to the cage of their former cage mates throughout behavioral neurophysiology data collection. Rats were maintained on a reverse light cycle (light: 8 p.m. – 8 a.m.), and all behavioral neurophysiology experiments were performed during the dark cycle. When necessary to encourage open field exploration, rats were temporarily placed on a food deprivation protocol. Five WT rats and one FXS rat experienced temporary food deprivation. With the provided food and treats consumed during open field exploration (see “*Open field behavioral task*” section below), rats on food deprivation maintained at least 97% of their free-feeding body weight.

After experiments had concluded, tissue (∼1-2 mm) was collected from anesthetized rats immediately prior to terminal perfusion. Tissue samples were then stored at −20°C and sent to Transnettyx Inc. to verify rats’ genotypes.

*Subjects for olfactory habituation/dishabituation testing.* Thirty-six rats (18 WT rats and 18 FXS rats) were included in the olfactory habituation/dishabituation behavioral task in this study. Two of the WT rats and four of the FXS rats were the same rats used for recording CA2 place cells. Rats who were included in both olfactory habituation/dishabituation tests and CA2 place cell recording studies completed olfactory testing prior to undergoing implantation surgery and CA2 data collection. All olfactory habituation/dishabituation testing was performed during the dark cycle. As with rats used for CA2 place cell recordings, tissue samples were collected from anesthetized rats immediately prior to euthanasia. Tissue samples were then stored at −20°C and sent to Transnettyx Inc. to verify rats’ genotypes.

*Subjects for analysis of oxytocin receptor expression.* Tissue was collected from three WT rats and nine FXS rats for oxytocin receptor immunostaining. Experimenters were blinded to rat genotype during tissue collection, immunostaining, imaging, and image analysis. Tissue samples were collected from anesthetized rats immediately prior to terminal perfusion. Tissue samples were then sent to Transnettyx Inc. for genotyping to verify rats’ genotyping. Following genotype verification, it was discovered that one cage had been assigned with the incorrect genotype. Estimates of oxytocin receptor levels for this cage were subsequently reassigned to the other genotype group. The mislabeled cage was a cage of FXS rats, which is why the FXS group had more rats than the WT group.

*Subjects for in vitro recordings.* CA2 intracellular recordings were made in acute hippocampal slices from four WT and four FXS male rats. Rats had free access to food and water and were housed on a reverse light-dark cycle (12 hours on/ 12 hours off). Rats were 4–5 months old at the time of slice preparation.

### Surgery and tetrode positioning for CA2 place cell recordings

“Hyperdrive” recording devices containing 14-21 independently movable tetrodes were implanted above the right dorsal hippocampus (anterior-posterior −3.8 mm from bregma, medial-lateral −3.0 mm from bregma). Tetrodes were constructed from 17 µm polymide-coated platinum-iridium (90/10%) wire (California Fine Wire, Grover Beach, California). The tips of the tetrodes designated for single unit recording were platinum-plated to reduce single-channel impedances to ∼150-300 kOhms. During surgery, eleven bone screws were placed in the skull and covered with dental acrylic to anchor the hyperdrive to the skull. Two of the screws were connected to the recording drive ground.

On the day of surgery, all tetrodes were lowered ∼1 mm into the brain. Tetrodes were then slowly lowered to the hippocampal pyramidal cell body layer over several weeks except for one tetrode that was designated as a reference electrode for differential recording. This tetrode was placed in an electrically quiet area of cortex at least 1 mm above the hippocampal cell body layer. The reference signal was duplicated using an MDR-50 breakout board (Neuralynx, Bozeman, Montana) and continuously recorded to ensure that unit activity or volume conducted hippocampal signals of interest (e.g., theta rhythms, sharp wave-ripples) were not detected.

Another tetrode was placed in the apical dendritic layer of CA1 to monitor hippocampal local field potential (LFP) signals; this tetrode helped to guide placement of the other tetrodes using estimated depth and electrophysiological hallmarks of the hippocampus (e.g., sharp wave-ripples).

### Data acquisition for CA2 place cell experiments

Data were acquired using a Digital Lynx system and Cheetah recording software (Neuralynx, Bozeman, Montana). The recording setup has been described in detail previously (Zheng et al., 2016). Briefly, LFP signals from one randomly chosen channel within each tetrode were continuously recorded at a 2000 Hz sampling rate and filtered in the 0.1–500 Hz band. Input amplitude ranges were adjusted before each recording session to maximize resolution without signal saturation. Input ranges for LFPs generally fell within ±2,000 to ±3,000 μV. To detect unit activity, all four channels within each tetrode were bandpass filtered from 600 to 6,000 Hz.

Putative spikes were detected when the filtered continuous signal on one or more of the channels exceeded 55 µV. Detected events were acquired with a 32,000 Hz sampling rate for 1 ms. For both LFPs and unit activity, signals were recorded differentially against a dedicated reference channel (see “*Surgery and tetrode positioning for CA2 place cell recordings*” section above).

Videos of rats’ behavior were recorded through the Neuralynx system with a resolution of 720 × 480 pixels and a frame rate of 29.97 frames/s. Rat position and head direction were tracked via an array of red and green or red and blue light-emitting diodes (LEDs) on a HS-54 or HS-27 headstage (Neuralynx, Bozeman, Montana), respectively.

### Open field behavioral task

Rats were familiarized to an open field arena (1 m x 1 m with 0.5 m wall height) for a minimum of three days before data collection began. Rats freely explored the open field arena for four 20-minute sessions per day, with 10-minute rest sessions preceding and following each active exploration session. During active exploration, small pieces of sweetened cereal or cookies were randomly scattered to encourage rats to explore the entirety of the arena. Rats had to cover at least 60% of the arena across each of the four sessions for a day to be included for further analysis. During each rest session, rats rested in a towel-lined, elevated flowerpot outside of the arena. A plexiglass standard rat housing cage was placed in one corner of the arena for all recording sessions. In the first and fourth sessions (A and A’), this cage contained only clean bedding. In the middle two sessions (B and B’) of the different experimental conditions, the cage contained social stimuli (Figure 1). In the Odor condition, a cage that had formerly housed familiar rat(s) and contained soiled bedding was used. In the Visual + Odor condition, a familiar rat was presented in its home cage containing soiled bedding. In the Control condition, a clean cage containing only clean bedding was presented in all four sessions.

**Figure 1.**
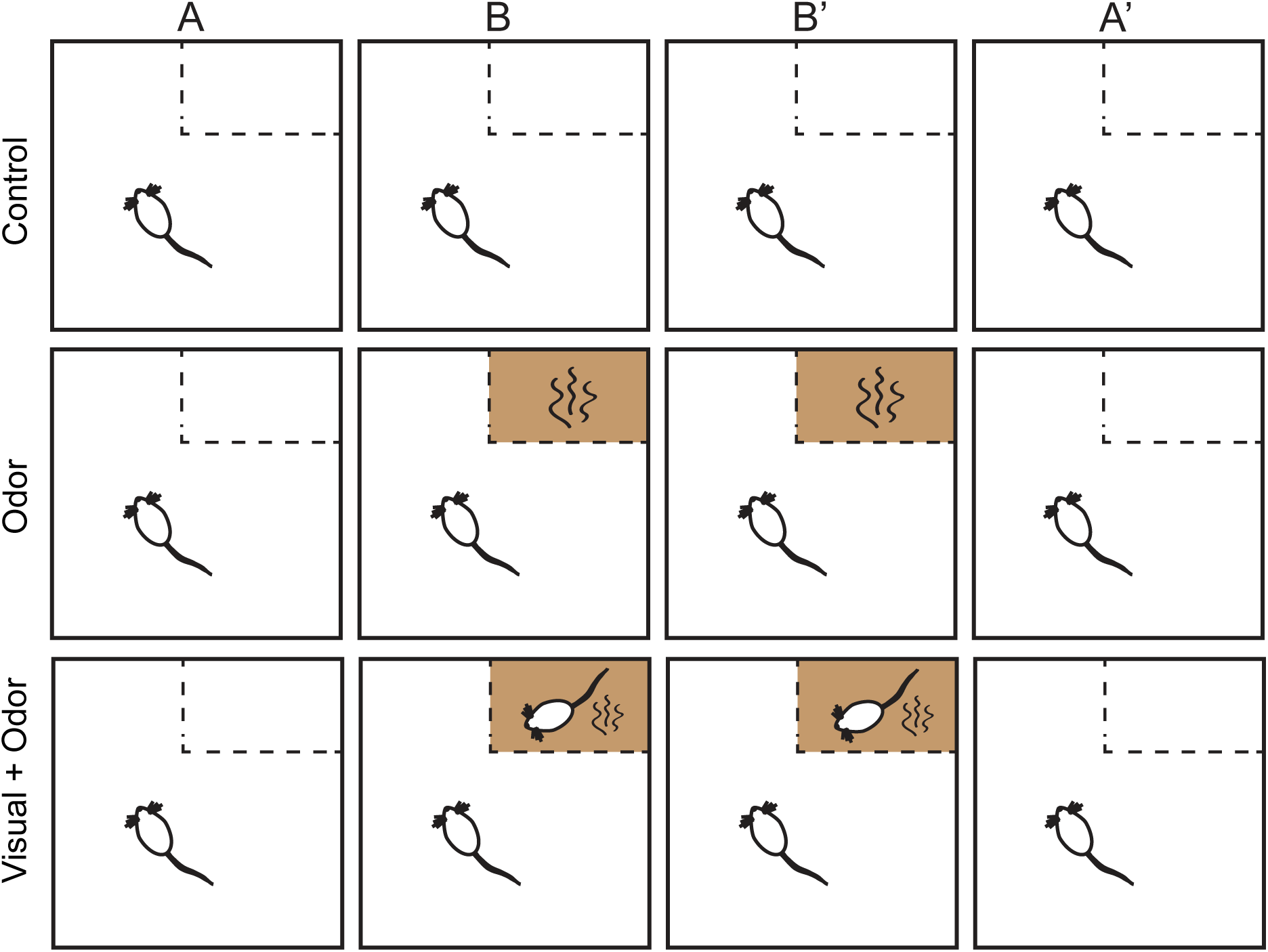
Behavioral task. Rats explored a 1-meter x 1-meter open field arena across four 20-minute sessions per day (Sessions A, B, B’, A’). A stimulus cage, which was a standard rat housing cage, was placed in a corner of the arena. In the first and final sessions (A and A’) of both experimental conditions (i.e., Odor and Visual + Odor) and in all four sessions of the Control condition, a clean cage containing clean bedding was presented. In the middle two sessions (B and B’) of the experimental conditions, a cage containing social stimuli was presented. In the Odor condition, the soiled bedding of a familiar rat or rats served as the social stimuli. In the Visual + Odor condition, a cage containing a familiar rat and the rat’s soiled bedding was presented.

### Histology for tetrode localization for CA2 place cell recordings

After place cell recordings were completed, rats were perfused with 4% paraformaldehyde solution in phosphate-buffered saline. Brains were cut coronally in 30 μm sections using a cryostat. For four of the rats, brain slices were stained with CA2-expressed protein striatum-enriched protein-tyrosine phosphatase (STEP; Cell Signaling, 4817, 1:1000 dilution, raised in mouse). Slices were then incubated in an anti-mouse secondary antibody and developed with 3,3-diaminobenzidine (brown) (as in Alexander et al., 2016). Slices were then counter-stained with cresyl-violet (purple/blue) to indicate cell bodies. Both the presence of STEP (Figure S1A) and the enlargement of cell bodies indicative of the CA2 region of the hippocampus (Lu et al., 2015) (Figure S1B) were used to identify CA2. In eight of the rats, brains were immunostained for CA2 marker Purkinje Cell Protein 4 (PCP4; Sigma-Aldrich Cat# HPA005792, 1:200 dilution, raised in rabbit) (Figure S1C) in lieu of STEP staining. Slides were then mounted using DAPI Fluoromount-G (Southern Biotech) to visualize cell bodies. Tetrode recording sites were identified by comparing locations across adjacent sections.

### Spike sorting and unit selection for CA2 place cell analyses

Spike sorting was performed manually using graphical cluster-cutting software (MClust, A.D. Redish, University of Minnesota, Minneapolis, Minnesota) run in MATLAB (Mathworks). Spikes were sorted using two-dimensional representations of waveform properties (i.e., energy, peak, and peak-to-valley difference) from four channels. A single unit was accepted for further analysis if the associated cluster was well isolated from, and did not share spikes with, other clusters on the same tetrode. Units were also required to have a minimum 1 ms refractory period. Units with mean firing rates above 5 Hz were considered putative interneurons and excluded from further analysis. In order to be considered active in the arena, a unit had to reach a peak firing rate of at least 1 Hz. In order to be included in the sharp wave-ripple firing analysis, a unit had to have valid clusters in both the active exploration and rest sessions. CA2 cell yields in each condition are reported for WT rats in Table 1 and for FXS in Table 2.

**Table 1.** CA2 place cell yields for WT rats.

| Genotype | Rat ID | Control | Odor | Visual<br>+ Odor |
| --- | --- | --- | --- | --- |
| WT | Rat 117 ("Lauren") | 7 | 5 | 0 |
| WT | Rat 122 ("Chase") | 51 | 32 | 0 |
| WT | Rat 165 ("Gus") | 34 | 56 | 26 |
| WT | Rat 256 ("Viggo") | 28 | 13 | 13 |
| WT | Rat 391 ("Daffodil") | 15 | 4 | 5 |
| WT | Rat 418 ("Hugo") | 20 | 35 | 41 |
|  | Total | 155 | 145 | 85 |

**Table 2:** CA2 place cell yields for FXS rats.

| Genotype | Rat ID | Control | Odor | Visual<br>+ Odor |
| --- | --- | --- | --- | --- |
| FXS | Rat 120 ("Brock") | 19 | 13 | 0 |
| FXS | Rat 125 ("JR") | 2 | 3 | 0 |
| FXS | Rat 330 ("Aires") | 0 | 1 | 1 |
| FXS | Rat 395 ("Elijah") | 6 | 1 | 4 |
| FXS | Rat 445 ("Paddy") | 7 | 7 | 5 |
| FXS | Rat 442 ("Pippin") | 33 | 35 | 26 |
|  | Total | 72 | 60 | 40 |

### Place cell remapping analyses

Methods used to create firing rate maps for each single unit were based on established methods from the place cell field that were used in our lab’s prior studies (Alexander et al., 2016; Robson et al., 2025; example rate maps for the present study shown in Figure 2). First, the arena was divided into 4 cm^2^ bins. The number of spikes that occurred within each bin was divided by the time spent in that bin to determine the firing rate. Only spikes that occurred while the rat was traveling 5 cm/s or faster were included. The rate map was smoothed with a two-dimensional Gaussian kernel (standard deviation = 6 cm). To determine if a place cell remapped during presentation of a social stimulus, a Pearson correlation coefficient *R* was calculated for each unit between pairs of rate maps from control and social stimuli sessions (i.e., A-B, B-B’, B’-A’, A-B’, and A-A’ session pairs, see Figure 1). To determine if spatial correlation coefficients differed between WT and FXS rats, or across session pairs (e.g., A-B and A-A’) or conditions (i.e., Odor, Visual + Odor, and Control), we used a generalized linear mixed model statistical analysis (IBM SPSS Statistics, version 29.0.2.0; see “*Experimental design and statistical analysis*” section below). The estimated mean spatial correlations for each genotype and condition, collapsed across session pairs, are shown in Figure 3A. Individual spatial correlation values for each cell are shown for each session pair and each condition in Figure 3B-F.

**Figure 2.**
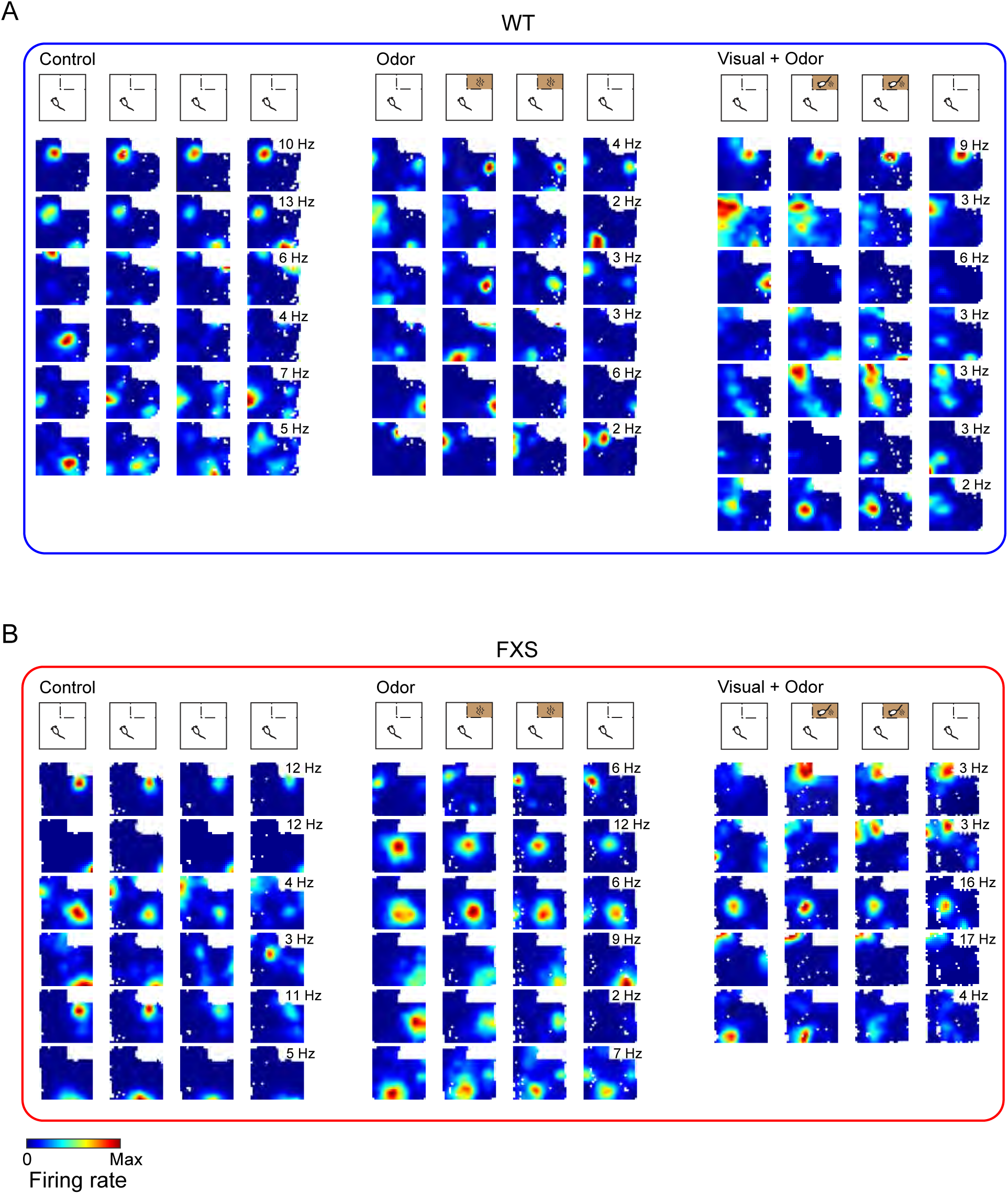
CA2 place cells in WT and FXS rats. Color-coded firing rate maps are shown for all place cells recorded on a single example tetrode across all conditions for WT (A) and FXS (B) rats. Rate maps are shown scaled to the maximal firing rate (shown inset) of each cell across all sessions. White pixels indicate places that were not visited during a session.

**Figure 3.**
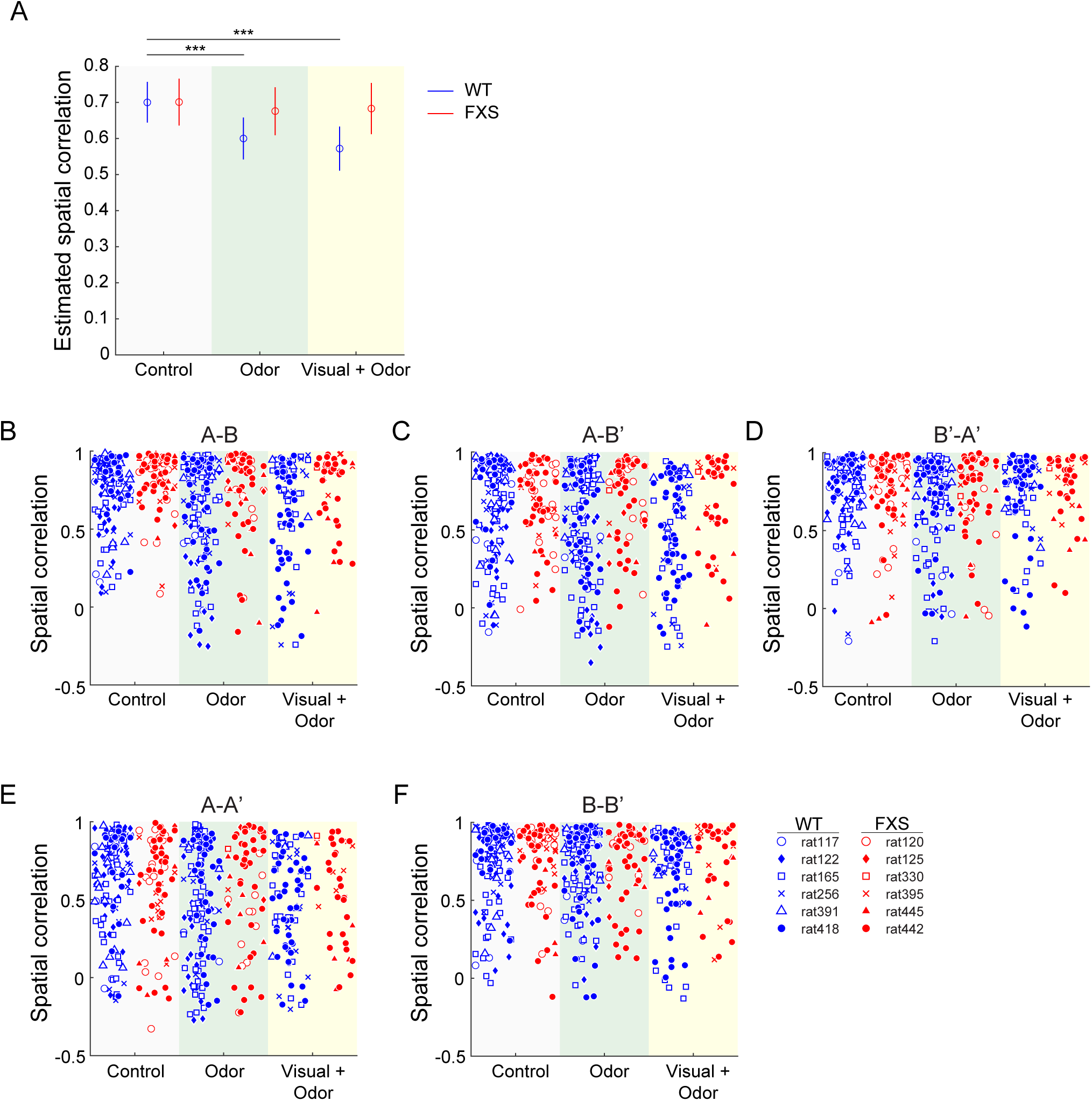
CA2 place cells changed their firing fields in response to social stimuli in WT, but not FXS, rats. (A) The estimated means of spatial correlation coefficients from the generalized linear mixed model are shown for each genotype across each condition for all session pairs combined. Dots represent estimated mean values, and error bars represent the estimated 95% confidence intervals. Spatial correlation coefficients were lower in the Odor and Visual + Odor conditions compared to the Control condition for WT rats. Spatial correlation coefficients did not differ across social and control conditions in FXS rats. (B-F) Spatial correlation measures are shown for the entire sample of CA2 place cells for all pairs of sessions across all conditions. Each marker represents a spatial correlation value for an individual place cell. Different symbols are used for CA2 place cells recorded from different rats. Data from WT and FXS rats are shown with blue and red symbols, respectively.

To determine if rate remapping occurred in CA2 place cells across genotypes (i.e., WT and FXS rats) and conditions (i.e., Odor, Visual + Odor, and Control), we calculated the rate overlap between pairs of sessions for each place cell. The rate overlap was defined as the ratio of the mean place field firing rate of the cell in each session, with the smaller of the two firing rates as the numerator (as in Robson et al., 2025). This analysis allowed us to determine if CA2 place cells changed their firing rates within a place field and therefore excluded cells that did not have an identified place field from a given session. This allowed us to compare firing rates without the confound of including cells that may gain or lose a field between sessions, which is defined as global remapping, not rate remapping. To determine if rate overlap differed across genotypes (i.e., WT and FXS rats), session pairs (i.e., A-B, A-B’, B’-A’, A-A’, B-B’), and conditions (i.e., Odor, Visual + Odor, and Control), we used a generalized linear mixed model statistical analysis (IBM SPSS Statistics, version 29.0.2.0; see “*Experimental design and statistical analysis*” section below). The estimated mean rate overlap values for each genotype and condition, collapsed across session pairs, are shown in Figure S2A. Individual rate overlap values for each cell are shown for each session pair and each condition in Figure S2B-F.

### Analyses of place cell properties

Spatial information (Figure 4A) was calculated as previously described (Skaggs et al., 1996) using the following formula:

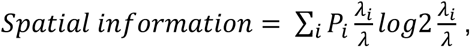

where *i* = the spatial bin index, *P_i_* = the probability of the rat being in the *i*th bin, *λ_i_* = the mean firing rate in the *i*th bin, and *λ* = the mean firing rate of the cell.

**Figure 4.**
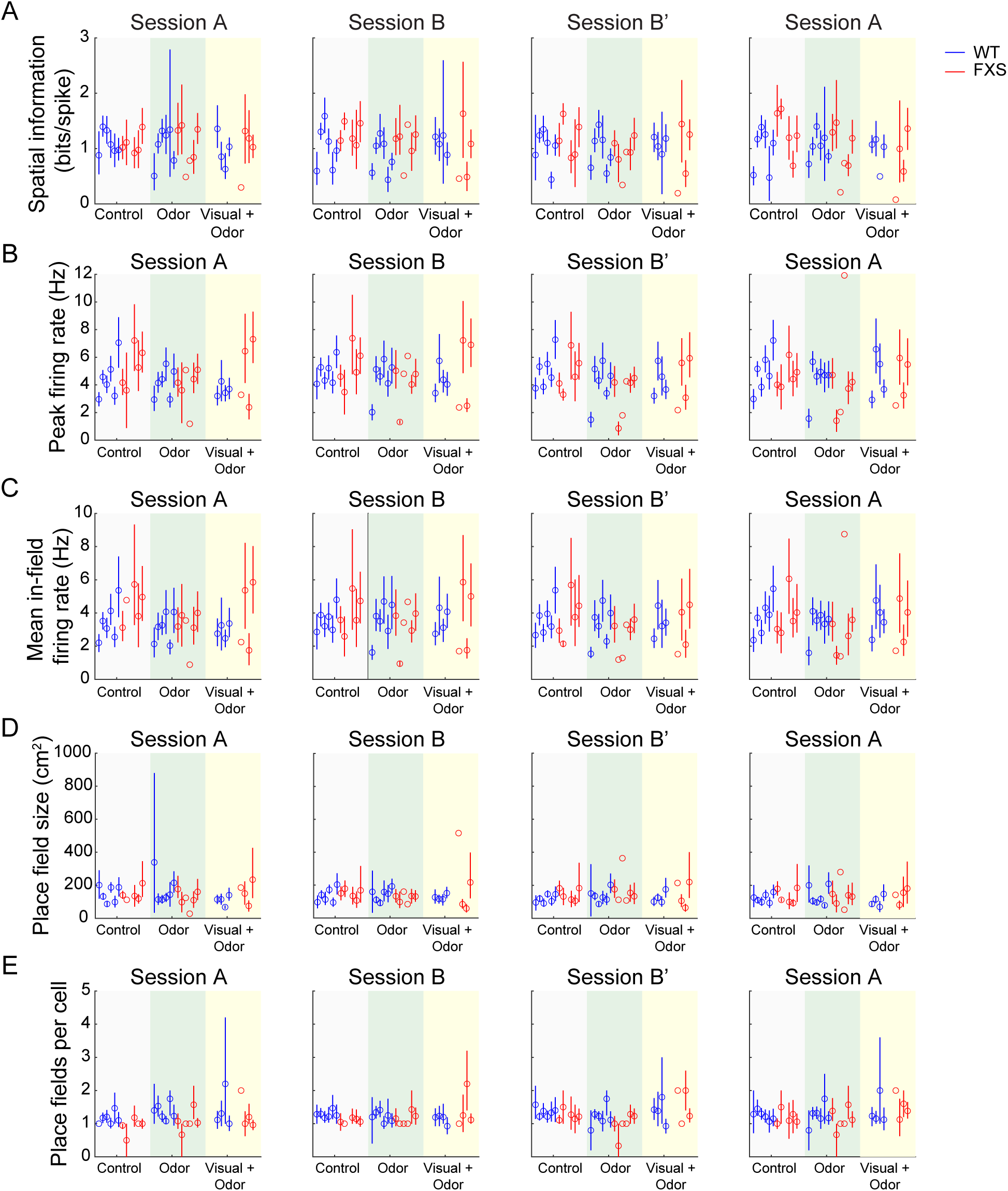
Basic firing and spatial coding properties of CA2 place cells did not differ between WT and FXS rats. For all plots, individual dots represent the mean value for each rat, and error bars represent 95% confidence intervals across all place cells within a rat. Data from WT rats and FXS rats are shown in blue and red, respectively. (A) Spatial information did not differ between WT and FXS rats across sessions and conditions. (B-C) CA2 place cells in FXS rats showed normal firing rates. Peak firing rates (B) and average in-field firing rates (C) of CA2 place cells did not differ between WT and FXS rats across sessions and conditions. (D) CA2 place field sizes were similar between genotypes and across sessions and experimental conditions. (E) The average number of place fields per place cell did not differ between genotypes across all sessions and conditions.

Place fields were separately identified for each cell in each session. Potential place fields were identified as contiguous bins of the smoothed rate map (see “*Place cell remapping analyses*” section above) that exceeded 50% of the peak firing rate in that session. In order to be included for further analysis, the peak firing rate of a place field had to exceed 1 Hz, and the size of the place field had to exceed 10 cm^2^.

### Exploration time analysis for open field behavioral task

Exploration of the stimulus cage was quantified as previously described (Robson et al., 2025). The arena was divided into 4 cm^2^ bins, and the time spent in each bin during the first two minutes of every session was determined for each day. These exploration maps were then smoothed with a two-dimensional Gaussian kernel (standard deviation = 4 cm). Maps were averaged within and then across rats within each genotype and plotted as a heat map for each condition (Figure 5A-B). The time spent within 12 cm of the cage was then calculated (Figure 5C).

**Figure 5.**
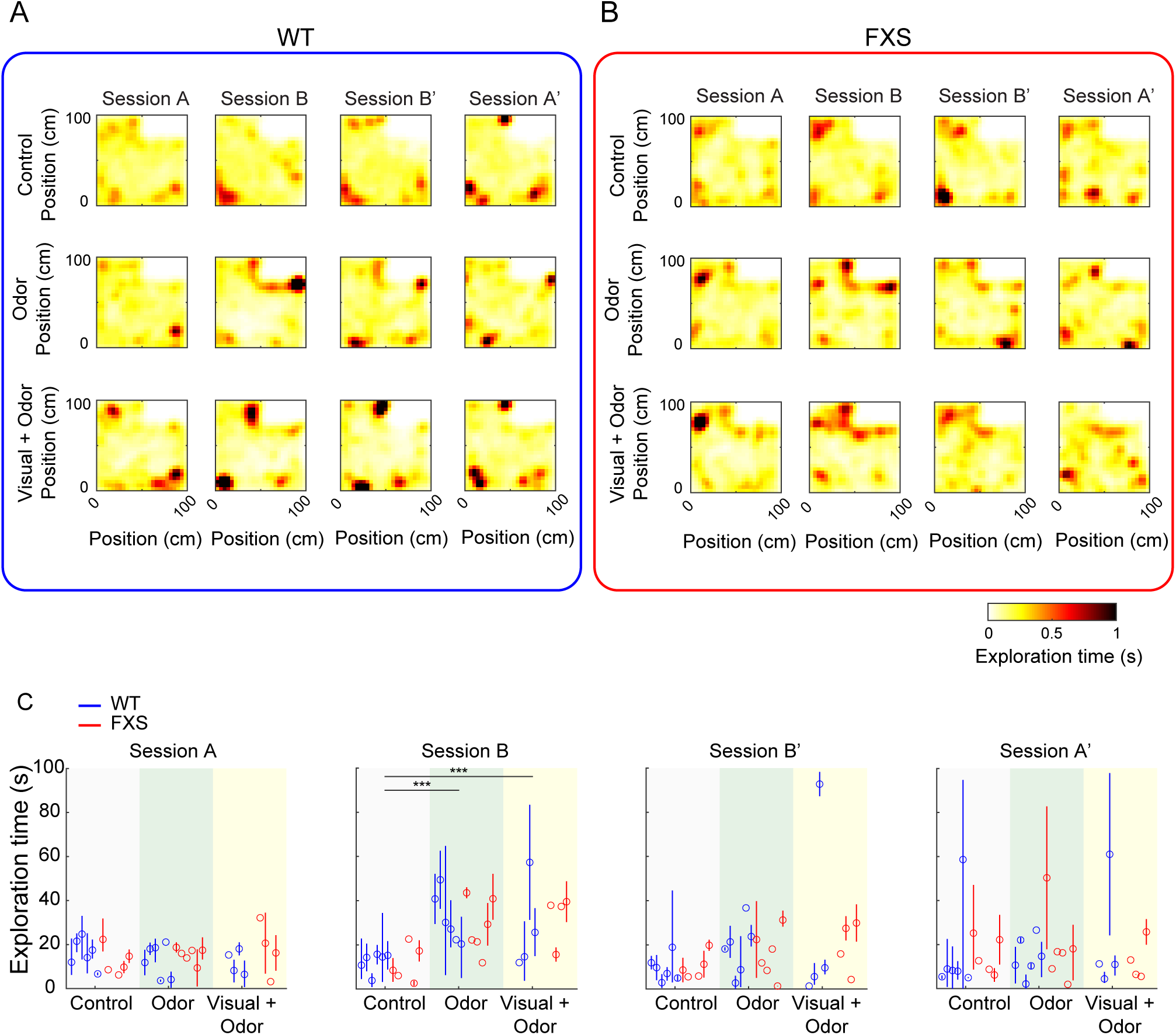
WT and FXS rats explored social stimuli similarly. (A-B) Heat maps show mean exploration times across locations in the arena, with the stimulus cage shown in the top right corner. Exploration times were calculated for the first 2 minutes of each individual session, averaged within each rat, and then averaged across rats within each genotype group for WT (A) and FXS (B) rats. (C) Time spent exploring locations close to the cage (within 12 cm) was calculated for each session and then averaged within a rat. Individual dots represent the mean exploration time for each rat, and error bars represent 95% confidence intervals across all sessions within a rat. Exploration times are shown in blue for WT rats and in red for FXS rats. Both WT and FXS rats increased their exploration time of the stimulus cage in session B for the Odor and Visual + Odor conditions but not the Control condition.

### Olfactory habituation/dishabituation task

To determine if rats could detect and distinguish between social odors, we used an olfactory habituation/dishabituation task (Yang & Crawley, 2009). First, rats were pre-acclimatized to the test cage containing a clean dry cotton tip applicator for 15 mins to reduce novelty-induced exploratory activity during the test. During the testing phase, rats were initially exposed to a cotton-tip applicator with no odor for 2-3 trials (“null” trials; N1-N3 in Figure 6). This was used to establish a baseline level of exploration for the object (i.e., the cotton-tip applicator). Following the null trials, rats were exposed to cotton tips with an unfamiliar social odor for four consecutive trials (“habituation” trials, H1-H4 in Figure 6). Following the habituation trials, rats were exposed to a cotton tip with odors from a second unfamiliar social odor (“dishabituation” trial, D in Figure 6). All trials were 90 seconds in length. Social odors were prepared by swiping soiled home cages of unfamiliar rats with cotton tip applicators. All trials were video recorded. Following completion of the experiment, the time rats spent sniffing the cotton tip (within ∼2 cm of the cotton tip and nose pointing towards the cotton tip) during each trial was scored by two independent scorers. The time spent sniffing was then averaged across the scorers. To account for differences in baseline activity levels across rats, we calculated normalized difference scores for each rat (*r*) and trial (*t*) using the following formula:

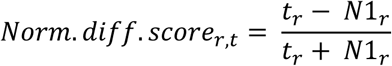

where *t_r_* was the sniffing time from the trial of interest for a given rat and *N1_r_* was the sniffing time from the first null trial for that rat.

**Figure 6.**
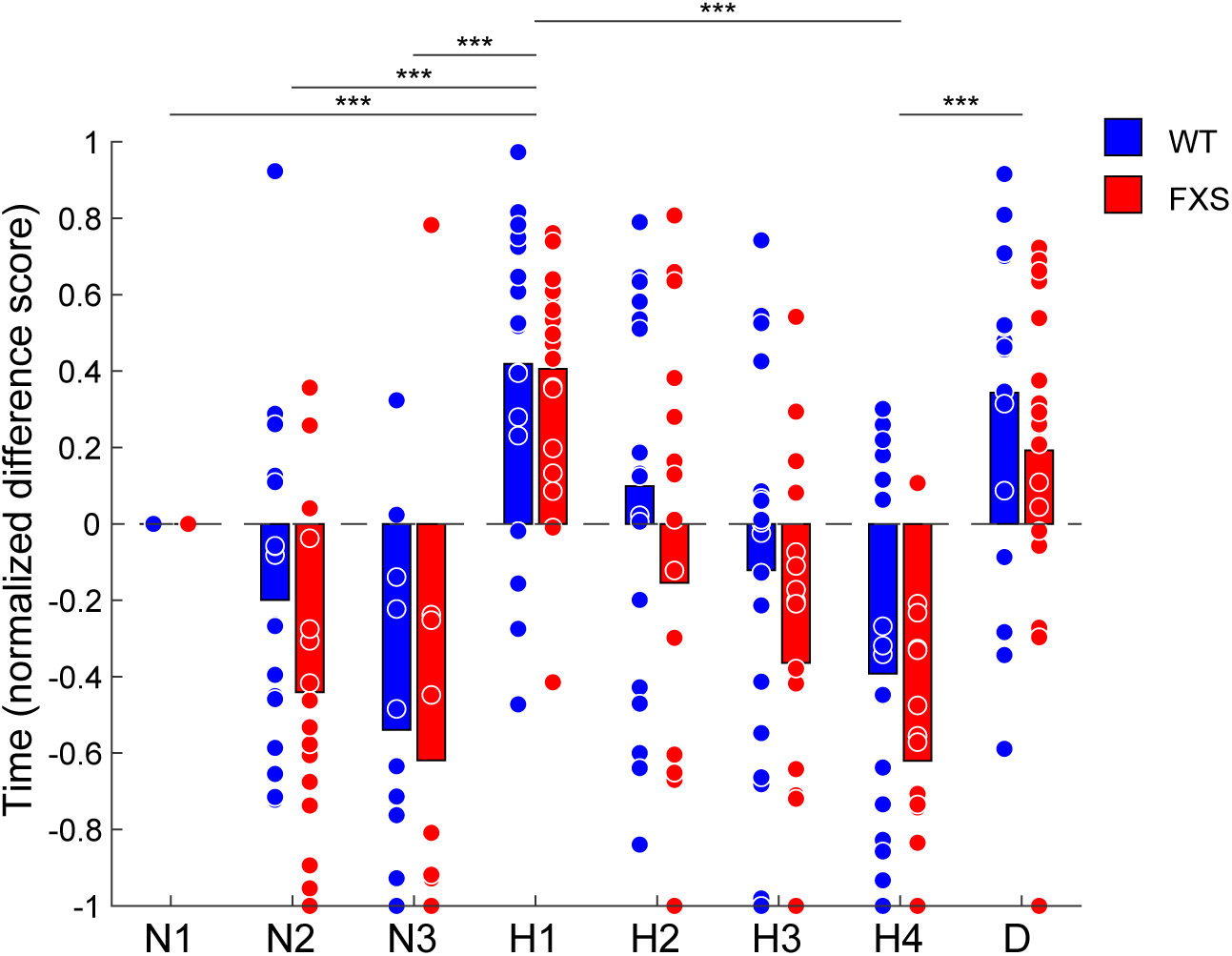
FXS rats could detect and differentiate between social odors. The normalized times spent sniffing a cotton-tip applicator is shown for trials with no odor (“null” trials, N1-N3), an initially unfamiliar social odor (“habituation” trials, H1-H4), and a second unfamiliar social odor (“dishabituation” trial, D). Bars indicate mean values within genotype and trial and dots indicate values from individual rats. Both WT (blue) and FXS (red) rats increased the time spent sniffing between null trial 3 (N3) and habituation 1 (H1), indicating that both WT and FXS rats successfully detected the presentation of a social odor. Both WT and FXS rats decreased the time spent sniffing across habituation trials (H1-4), indicating that FXS rats habituated normally to repeated presentations of the same odor. Both WT and FXS rats increased time spent sniffing between habituation trial 4 (H4) and the dishabituation trial (D), indicating that both WT and FXS rats distinguished between different social odors.

### Immunohistostaining for oxytocin receptors

Rats (three WT rats and nine FXS rats, aged 25 weeks) were perfused in phosphate-buffered saline (PBS) followed by 4% paraformaldehyde solution in PBS. Brains were stored overnight in 4% paraformaldehyde solution in PBS. On the day after perfusion, brains were cut coronally in 30 μm sections using a cryostat and mounted onto slides. Sections were kept frozen at −80 degrees Celsius until tissue from all rats had been collected. Immunostaining was then performed simultaneously on all sections from both genotypes. First, all slides were defrosted to room temperature and washed multiple times with water, boiling water, and Tris-buffered saline (TBS) + 0.3% Triton X-100. Sections were then blocked for 2 hours with 5% Normal Goat Serum (NGS). Sections were then incubated overnight with the following primary antibodies diluted in blocking buffer: Mouse anti-RGS14 (1:1000; Antibodies Inc) and rabbit anti-oxytocin receptor (1 μg/ml; Moses V. Chao, Robert C. Froemke, NYU School of Medicine Cat# OXTR-2 antibody, RRID:AB_2571555). The following day, slides were washed with Tris-buffered saline with Tween-20 (TBST) and then incubated for two hours with the following secondary antibodies: goat anti-mouse ALEXA 555, A-21422 (1:500; Thermo Fisher) and goat anti-rabbit ALEXA 488, A-11008 (1:500, Thermo Fisher). Slides were then washed with TBST and TBS. Lastly, slides were mounted using DAPI Fluoromount-G (Southern Biotech).

### Quantification of OXTR-2 expression

Slides were imaged using a Zeiss AXIO Zoom.V16 (software Zen 3.11 Pro) microscope with an Axiocam512 color camera with filters for imaging DAPI (blue, excitation range 350-360 nm, emission range 460nm), Cy3 (orange-red excitation range 513-556, emission 570-613), and

EGFP (Enhanced Green Fluorescent Protein, excitation about 470, emission about 510) at a magnification of 77x. For imaging sections that were immunostained with multiple antibodies, a standardized exposure time was chosen for each filter: DAPI 200 ms, Cy3 350 ms, and EGFP 280 ms. The levels of oxytocin receptors in each section were estimated via the following procedures. First, blinded experimenters took photomicrographs of WT and FXS sections using the same microscope settings. Images were saved and subsequently analyzed by blinded experimenters using Image J software (ImageJ 1.54g). Images taken with the blue (DAPI), green (OXTR-2), and red (RGS14) settings were opened and calibrated to the same scale.

Regions of interest (i.e., CA3, CA2, and CA1) were defined in all 3 sets of images using the RGS14 staining as a marker for CA2. To estimate OXTR-2 levels, background fluorescence was removed, and estimates were obtained for mean fluorescence (average pixel intensity within each region of interest) and total (integrated) fluorescence. To determine if OXTR-2 expression differed between WT and FXS rats, or across hippocampal subregions, we used a generalized linear mixed model statistical analysis (IBM SPSS Statistics, version 29.0.2.0; see “*Experimental design and statistical analysis*” section below). The estimated corrected fluorescence from the generalized linear mixed model for each genotype and subregion is shown in Figure 7C.

**Figure 7.**
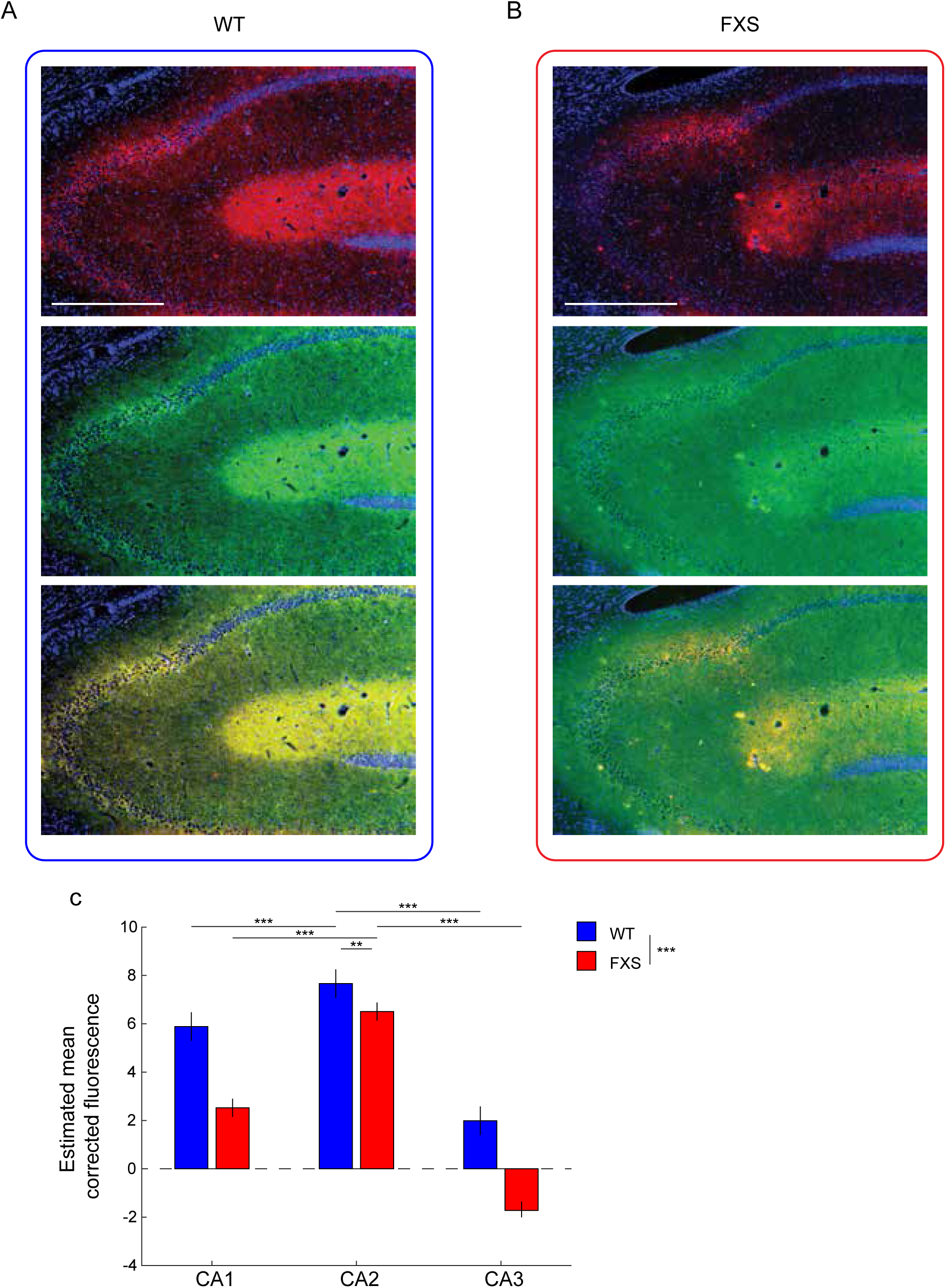
Low oxytocin receptor expression levels in CA2 of FXS rats. (A-B) Example images of CA2 immunostaining in WT (A) and FXS (B) rats for a CA2 marker (Regulator of G-protein Signaling 14 (RGS14), red, top row) and an oxytocin receptor antibody (OXTR-2, green, middle row). DAPI nuclear staining is shown in blue. Merged images for RGS14 and OXTR-2 immunostaining are shown in the bottom row. Scale bar: 500 µm. (C) Plots show fluorescence estimates (mean ± 95% confidence intervals) across hippocampal subregions for WT (blue) and FXS (red) rats. Oxytocin receptor expression across subregions was lower in FXS rats than in WT rats.

### Slice preparation for in vitro recording

Rats were anesthetized using a ketamine/xylazine cocktail (100/10 mg/kg) and then underwent cardiac perfusions with ice-cold saline consisting of (in mM): 2.5 KCl, 1.25 NaH_2_PO_4_, 25 NaHCO_3_, 0.5 CaCl_2_, 7 MgCl_2_, 7 dextrose, 205 sucrose, 1.3 ascorbic acid, and 3 sodium pyruvate (bubbled constantly with 95% O_2_/5% CO_2_ to maintain pH at ∼7.4). Following perfusion, the brain was removed and sliced into 300 µM parasagittal sections from the middle hippocampus using a vibrating tissue slicer (Vibratome 300, Vibratome Inc). The slices were placed in a chamber filled with artificial cerebral spinal fluid (aCSF) consisting of (in mM): 125 NaCl, 2.5 KCl, 1.25 NaH_2_PO_4_, 25 NaHCO_3_, 2 CaCl_2_, 2 MgCl_2_, 10 dextrose, 1.3 ascorbic acid and 3 sodium pyruvate (bubbled constantly with 95% O_2_/5% CO_2_) for 30 minutes at 35°C and then held at room temperature until time of recording.

### In vitro electrophysiology

Slices were placed in a submerged, heated (33-34 C°) recording chamber and continually perfused with aCSF (in mM): 125 NaCl, 3 KCl, 1.25 NaH_2_PO_4_, 25 NaHCO_3_, 2 CaCl_2_, 1 MgCl_2_, 10 dextrose, and 3 sodium pyruvate (bubbled constantly with 95% O_2_/5% CO_2_). Ionotropic glutamatergic and GABAergic synaptic transmission were blocked with 20 µM DNQX, 25 µM D-AP5, and 2 µM gabazine. Pyramidal neurons within the CA2 region were visualized with a Zeiss AxioScope under 60x magnification.

Current clamp recordings were made using a Dagan BVC-700 or Sutter IPA amplifier and Sutterpatch acquisition software. Data were sampled at 20-50 kHz, filtered at 3 kHz, and then digitized by a Dendrite interface or integrated A/D board in the Sutter IPA (Sutter). The internal recording solution consisted of (in mM): 135 K-gluconate, 10 HEPES, 7 NaCl, 7 K_2_-phosphocreatine, 0.3 Na-GTP, 4 Mg-ATP (pH corrected to 7.3 with KOH). Recording electrodes were pulled from borosilicate glass and had an open tip resistance of 4-6 MΩ. Series resistance was compensated using the bridge balance circuit and was monitored throughout the experiment. Recordings in which the series resistance exceeded 35 MΩ or increased by more than 30% were excluded. The resting membrane potential was noted immediately after establishing the whole cell recording configuration.

### Drugs for in vitro electrophysiology experiments

DNQX, APV, and gabazine were purchased from Hello Bio. Thr^4^-Gly^7^ oxytocin (TGOT) was purchased from Bachem. All drugs were prepared from a 1000x stock solution in water.

### Experimental design and statistical analyses

Whenever possible, experimenters were blinded to genotype while collecting and analyzing place cell data (i.e., during spike sorting). A period of limited availability of FXS rats from the supplier resulted in unblinded collection and analysis of data from three rats (rats 418, 442, and 445). All video scoring for the olfactory habituation/dishabituation task was performed by scorers blinded to genotype. Experimenters were blinded to genotype during OXTR-2 immunostaining and quantification. Two experimenters conducted the in vitro electrophysiology experiments.

One was blinded to genotype, but the other was not.

We used a generalized linear mixed model approach (IBM SPSS version 29.0.2.0) to assess differences between genotypes (i.e., WT and FXS rats) for in vivo neurophysiology, behavior, and histology data. For all analyses, rats were set as the subjects, and genotype was included as a fixed factor. Post-hoc tests were performed if a significant interaction or main effect was observed and used the Bonferroni correction. For place cell analyses, multiple place cells were nested within rats. To determine if spatial correlation coefficients (Figure 3) or rate overlap values (Figure S2) differed across genotypes, conditions, and session pairs, condition and session pair were included as fixed factors, and session pairs were repeated measures within rats. A three-way interaction between genotype, condition, and session pair was included in the model, as was a two-way interaction between genotype and condition. To assess place cell properties (Figure 4), session was included as a repeated measure within rats. In addition to the main effect of genotype, the model included: a three-way interaction between genotype, condition, and session; a two-way interaction between genotype and session; and a two-way interaction between condition and session.

To verify that our statistical tests were sufficiently powered given that a lower number of CA2 place cells were recorded in FXS rats than in WT rats (see Tables 1 and 2), we performed the following analyses. First, we performed a simulation-based power analysis in which we randomly down-sampled the number of CA2 place cells from WT rats to match the number recorded in FXS rats for each condition (i.e., 72 cells in the Control condition, 60 cells in the Odor condition, and 40 cells in the Visual + Odor condition) 100 times. We then used a generalized linear mixed model to test for a statistically significant difference between the Control condition and the Odor and Visual + Odor conditions for each simulation. We included an interaction between condition and session and main effects of condition and session in the model. Post-hoc comparisons were made using the Bonferroni correction when a significant effect was found. We tallied the number of simulations in which the post-hoc pairwise comparison showed a significant difference between the control and experimental conditions. We did this separately for Control and Odor, and Control and Visual + Odor, comparisons to obtain power estimates for each. We additionally ran permutation tests to compare the effects of condition on spatial correlation values for each genotype and condition. The permutation tests shuffled condition 1,000 times to obtain a null distribution for each spatial correlation value.

Monte-Carlo p-values were calculated using the formula:

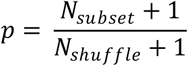

where *N_subset_* is the number of shuffles with values greater than or equal to the observed value (two-tailed test), and *N_shuffle_* is the total number of shuffles.

To compare social stimuli exploration times across genotypes (Figure 5), days were included as random factors nested within rats, as conditions were repeated on different days within rats.

Sessions were set as a repeated measure. In addition to the main effect of genotype, the model included a three-way interaction between genotype, condition, and session and a two-way interaction between condition and session.

To analyze olfactory stimuli investigation times in the habituation/dishabituation task (Figure 6), trial was set as a repeated measure. In addition to the main effect of genotype, a main effect of trial and a two-way interaction between genotype and trial was included in the model.

To compare the expression of oxytocin receptors across genotypes and hippocampal subregions (Figure 7), multiple fluorescent estimates from brain sections were nested within each subregion and set as repeated measures. In addition to genotype, subregion was included as a fixed factor in the model.

Statistical tests for in vitro electrophysiology data (Figure 8) were also performed in SPSS (IBM SPSS version 29.0.2.0). To compare the resting membrane potential of CA2 neurons across WT and FXS rats, a univariate analysis of variance was performed with genotype as the between-subjects factor. A general linear model was used to assess the effects of TGOT application on CA2 neurons’ resting membrane potential, with drug treatment (baseline vs. TGOT) as a repeated measure and genotype as a between-subjects factor.

**Figure 8.**
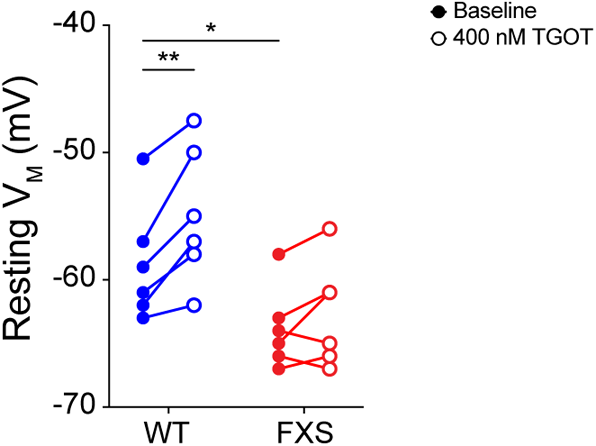
Oxytocin depolarized the resting membrane potential in CA2 neurons in WT rats but not FXS rats. Resting membrane potential (V_M_) measurements are shown during baseline and after 15 minutes of 400 nM TGOT application. Dots indicate data from individual cells. TGOT significantly depolarized the V_M_ of WT CA2 neurons (blue; n = 6 cells from 4 rats) but not FXS CA2 neurons (red; n = 6 cells from 4 rats).

### Data and code availability

The analysis code is available on GitHub (https://github.com/mmdonahue/FMR1CA2_PUBLICSHARE). Data will be made available upon reasonable request.

## Results

### CA2 place cells in FXS rats did not remap to social stimuli

Our prior studies have shown that CA2 place cells remap to social stimuli (Alexander et al., 2016; Robson et al., 2025) and, specifically, that this remapping is driven by the olfactory content of social stimuli (Robson et al., 2025). As FXS rats have atypical social behaviors (Hamilton et al., 2014; Tian et al., 2017; Wong et al., 2020), the primary goal of this study was to determine if CA2 place cells in FXS rats respond normally to the presentation of social stimuli.

Thus, we presented social stimuli to FXS and WT control rats while recording CA2 place cell activity. We compared CA2 place cell firing patterns between WT and FXS rats across social and control conditions (Figure 1).

We have previously shown that the presentation of a conspecific in a cage with soiled bedding or the presentation of a soiled cage alone induces global remapping, but not rate remapping, in a subset of CA2 place cells (Alexander et al., 2016; Robson et al., 2025). To test if similar remapping occurs in CA2 place cells of FXS rats, we compared CA2 place cell firing patterns in WT and FXS rats exploring an open field arena in which social stimuli (i.e., a familiar rat in a soiled cage or a soiled cage alone) were presented. CA2 place cell firing patterns in WT and FXS rats during presentation of social stimuli were compared to CA2 place cells recorded in a control condition in which no social stimuli were presented in the stimulus cage (Figure 1).

Our recordings showed seemingly normal place cells in CA2 of FXS rats (Figure 2). To quantify the extent to which CA2 place cells exhibited global remapping (i.e., a change in place cell firing patterns in which place field locations change) when social stimuli were presented, we calculated spatial correlation coefficients between rate maps from pairs of sessions (Figure 3). We found that spatial correlation coefficients were lower for the Odor and Visual + Odor conditions compared to the Control condition in WT rats (Figure 3; generalized linear mixed model, no interaction between genotype, condition, and session pair (F(20,2755) = 1.2, p = 0.244); significant interaction between genotype and condition (F(2,2755) = 6.9, p = 0.001); significant differences in post-hoc tests for WT rats for Control vs. Odor (t(2755) = 6.5, p < 0.001) and Control vs. Visual + Odor (t(2755) = 6.8, p < 0.001)). However, there were no differences between the Control condition and either of the social conditions in FXS rats (Figures 2B and 3; no differences in post-hoc tests for FXS rats for Control vs. Odor (t(2755) = 1.1, p = 0.809) and Control vs. Visual + Odor (t(2755) = 0.7, p = 0.973)). These results suggest that CA2 place cells in FXS rats do not remap when social stimuli, namely a familiar conspecific in a cage with social odors or social odors in the absence of a rat, are presented.

Because a lower number of place cells were recorded in FXS rats than in WT rats (see Tables 1 and 2), we performed analyses to determine whether our tests were sufficiently powered to detect CA2 place cell remapping (Alexander et al., 2016; Robson et al., 2025) using the effect size observed in WT rats. First, we used a simulation-based power analysis in which we the number of CA2 place cells from WT rats was randomly down-sampled 100 times to match the number recorded in FXS rats. We then used a generalized linear mixed model to test for a statistically significant difference between the Control condition and the Odor and Visual + Odor conditions for each simulation. We found that there was a significant main effect of condition and a significant post-hoc pairwise comparison between the Control and Odor conditions 87% of the time. There was a significant main effect of condition and a significant post-hoc pairwise comparison between the Control and Visual + Odor conditions 92% of the time. We additionally ran permutation tests to compare the effects of condition on spatial correlation values for each genotype and condition. We found that spatial correlation values were lower in both the Odor (permutation test, 1000 shuffles of condition, effect size = −0.36, p < 0.001) and Visual + Odor (permutation test, 1000 shuffles of condition, effect size = −0.41, p < 0.001) conditions than in the Control condition in WT rats. However, there were no significant differences between the Control condition and the Odor or Visual + Odor conditions in FXS rats (Control vs. Odor: permutation test, 1000 shuffles of condition, effect size = −0.09, p = 0.27; Control vs. Visual + Odor, permutation test, 1000 shuffles of condition, effect size = −0.06, p = 0.51). These results suggest that the lack of CA2 place cell remapping observed in FXS rats was not explained by the lower number of place cells recorded in FXS rats.

Although CA2 place cells in FXS rats did not globally remap to a social stimulus, it is possible that CA2 place cells respond to social stimuli by rate remapping. To test this possibility, we calculated the rate overlap between pairs of sessions for CA2 place cells in WT and FXS rats (Figure S2). We have previously reported that CA2 place cells did not rate remap to the presentation of social stimuli in WT rats (Robson et al., 2025). In the present study, we also did not observe rate remapping to social stimuli in CA2 place cells in FXS rats (generalized linear mixed model, no interaction between genotype, condition, and session pair (F(8,2718) = 0.9, p = 0.511), no interaction between genotype and condition (F(2,2718) = 0.8, p = 0.433), no main effect of genotype (F(1,2718) < 0.1, p = 0.901), no main effect of condition (F(2,2718) = 1.4, p = 0.252)).

### CA2 place cells exhibited normal firing properties in FXS rats

The lack of remapping to social stimuli observed in CA2 place cells in FXS rats raised a question of whether basic firing or spatial coding properties of CA2 place cells were abnormal in FXS rats. Thus, we next assessed whether baseline firing properties of CA2 place cells differed between WT and FXS rats. While CA1 place cell activity has been characterized in both FXS mouse and rat models (Arbab, Battaglia, et al., 2018; Arbab, Pennartz, et al., 2018; Asiminas et al., 2022; Donahue, Robson, & Colgin 2025; Talbot et al., 2018), basic firing and spatial coding properties of CA2 place cells in FXS rodent models have not previously been described.

In addition to quantifying remapping across sessions in which environmental conditions change, spatial correlation coefficients can also be calculated across sessions in an unchanging environment to quantify place cell stability. We calculated spatial correlations between session pairs in the Control condition and observed no significant differences between WT and FXS rats (Figure 3; generalized linear mixed model, no difference between WT and FXS in post hoc test for Control condition, t(2755) < 0.1, p = 0.987)). This suggests that the baseline stability of CA2 place cells in an unchanged environment is similar between WT and FXS rats.

We next examined firing properties of CA2 place cells in WT and FXS rats during individual sessions. We found that spatial information transmitted by CA2 place cells did not differ between WT and FXS rats across all conditions and sessions (Figure 4A; generalized linear mixed model, no interaction between genotype, condition, and session (F(8,2204) = 0.5, p = 0.863); no interaction between genotype and session (F(2,2204) = 0.3, p = 0.803); no main effect of genotype (F(1,2204) = 0.2, p = 0.674)). Average in-field firing rates (Figure 4B; generalized linear mixed model, no interaction between genotype, condition, and session (F(8,2016) = 1.1, p = 0.366); no interaction between genotype and session (F(3,2016) = 1.8, p = 0.138); no main effect of genotype (F(1,2016) = 0.2, p = 0.668)) and peak firing rates (Figure 4C; generalized linear mixed model, no interaction between genotype, condition, and session (F(8,2016) = 1.1, p = 0.340); no interaction between genotype and session (F(3,2016) = 1.9, p = 0.128); no main effect of genotype (F(1,2016) = 0.2, p = 0.673)) of CA2 place cells were also similar between WT and FXS rats. Also, CA2 place field size (Figure 4D; generalized linear mixed model, no interaction between genotype, condition, and session (F(8,2016) = 0.2, p = 0.277); no interaction between genotype and session (F(3,2016) = 0.3, p = 0.857); no main effect of genotype (F(1,2016) = 0.8, p = 0.380)) and average number of place fields per place cell per session (Figure 4E; generalized linear mixed model, no interaction between genotype, condition, and session (F(8,2204) = 0.9, p = 0.539); no interaction between genotype and session (F(3,2204) = 1.3, p = 0.263); no main effect of genotype (F(1,2204) = 0.7, p = 0.389)) did not differ between WT and FXS rats. Altogether, these results indicate that CA2 place cells represent spatial locations similarly in FXS and WT rats.

### FXS rats explored and perceived social stimuli normally

A possible explanation of why CA2 place cells failed to remap to social stimuli in FXS rats could be reduced exploration of, or attention to, social stimuli in FXS rats. To explore this possibility, we calculated the amount of time spent exploring the stimulus cage across conditions for WT rats and FXS rats (Figure 5). We first created heat maps to visualize the amount of time rats spent exploring locations across the open field arena and noted that both WT and FXS rats spent a high amount of time near the stimulus cage in the Odor and Visual + Odor conditions (Figure 5A-B). To quantify these observations, we calculated the time spent near the stimulus cage across conditions and sessions for all rats. We found that both WT and FXS groups of rats increased the time they spent near the stimulus cage in Session B during the Odor and Visual + Odor conditions (Figure 5C; generalized linear mixed model, no interaction between genotype, condition, and session (F(11,260) = 0.4, p = 0.935); significant interaction between condition and session (F(11,260) = 5.4, p < 0.001); significant differences in post-hoc tests for Session B for Control vs. Odor (t(260) = −5.3, p < 0.001) and Control vs. Visual + Odor (t(260) = −4.1, p < 0.001)). This indicates that WT and FXS rats explored social stimuli to a similar extent.

We have previously shown that the olfactory content of social stimuli is necessary for CA2 place cell remapping to social stimuli (Robson et al., 2025). Although structural and functional changes have been identified in the olfactory system of FXS rodent models (Bodaleo et al., 2019), these differences appear to impact olfactory sensitivity and fine odor discrimination, but not the detection of odors or the discrimination of monomolecular odors (Kuruppath et al., 2023; Nitenson et al., 2015). Still, it is possible that impaired olfactory perception of social stimuli explains the lack of CA2 place cell remapping to social stimuli in FXS rats. To assess whether olfactory perception and discrimination were impaired in FXS rats, we assessed olfactory function in FXS rats using an olfactory habituation/dishabituation task (Figure 6). This task tests rats’ ability to recognize an odor when it is initially presented, habituate to the presence of the same odor across repeated presentations (“habituation” trials), and distinguish between odors after the presentation of a novel odor in the last trial (“dishabituation” trial). We found that FXS rats could detect the presence of a social odor normally during the first habituation trial (Figure 6; generalized linear mixed model, no interaction between genotype and trial (F(7,259) = 0.6, p = 0.722); significant main effect of trial (F(1,259) = 126.9, p < 0.001); significant differences in post-hoc tests for Null 1 vs. Habituation 1 (t(259) = −4.2, p = 0.001), Null 2 vs. Habituation 1 (t(259) = −7.4, p < 0.001), Null 3 vs. Habituation 1 (t(259) = −9.8, p < 0.001)). Further, both WT and FXS rats decreased their sniffing of the same social odor across repeated presentations (significant difference in post-hoc test for Habituation 1 vs. Habituation 4 (t(259) = 9.2, p < 0.001)), indicating normal habituation to a social odor in FXS rats. FXS rats also increased their sniffing in the same manner as WT rats when a novel social odor was introduced during the dishabituation trial (significant difference in post-hoc test for Habituation 4 vs. Dishabituation (t(259) = −8.7, p < 0.001)). This indicates that FXS rats detected social odors, and distinguished between different social odors, in the same way as WT rats.

### Impaired oxytocin modulation of area CA2 in FXS rats

The neuropeptide oxytocin plays a key role in social behaviors and social memory (Froemke & Young 2022). High expression of oxytocin receptors is a characteristic feature of area CA2 (Mitre et al. 2016), and oxytocin receptors in CA2 are essential for social functions (Raam et al. 2017, Lin et al. 2018, Tsai et al. 2022). Thus, we hypothesized that reduced responses to social stimuli in CA2 place cells could be due to deficient oxytocin receptor expression in CA2 of FXS rats. To test this hypothesis, we immunostained hippocampal tissue from WT and FXS rats with OXTR-2, an antibody that is highly specific for oxytocin receptors (Mitre et al. 2016). We then compared expression of oxytocin receptors in area CA2, and neighboring subregions CA1 and CA3, in WT and FXS rats (Figure 7). Significant differences in the mean intensity of OXTR-2 immunostaining were observed across subregions and genotypes (Figure 7C; generalized linear mixed model, significant interaction between genotype and subregion (F(2,701) = 17.1, p < 0.001)). In WT rats, oxytocin receptor expression levels were higher in CA2 than in CA1 and CA3 (generalized linear mixed model, significant main effect of subregion for WT rats (F(2,174) = 93.7, p < 0.001) and significant differences in post-hoc tests for CA1 vs. CA2 (p < 0.001) and CA3 vs CA2 (p < 0.001)), as expected based on earlier results from WT mice (Mitre et al. 2016). Similarly, oxytocin receptor expression in FXS rats was higher in CA2 than in CA1 and CA3 (generalized linear mixed model, significant main effect of subregion for FXS rats (F(2,527) = 459.7, p < 0.001) and significant differences in pairwise comparisons for CA1 vs. CA2 (p < 0.001) and CA3 vs CA2 (p < 0.001)). We additionally found that oxytocin receptor expression levels were significantly lower in FXS rats than in WT rats across all subregions (generalized linear mixed model, significant main effect of genotype (F(1,701) = 160.9, p < 0.001)) and in CA2 tested individually (generalized linear mixed model, significant main effect of genotype (F(1,233) = 9.8, p = 0.002)). These results raise the possibility that reduced responses to oxytocin in CA2 neurons contribute to diminished responses to social stimuli observed in CA2 place cells in FXS rats.

Thus, we next tested whether CA2 neurons in FXS rats exhibit reduced responses to oxytocin receptor activation. Prior work in WT mice showed that CA2 neurons depolarize in response to oxytocin, and such depolarizing responses were proposed to promote social remapping in CA2 (Tirko et al. 2018). Accordingly, we hypothesized that CA2 neurons would depolarize in response to oxytocin administration in slices from WT rats but not FXS rats. To test this hypothesis, we performed whole-cell current clamp recordings of CA2 pyramidal neurons and tested how bath application of the specific oxytocin receptor agonist Thy^4^,Gly^7^-oxytocin (TGOT, 400 nM) affected the resting membrane potential. First, we found that the baseline resting membrane potential of CA2 pyramidal neurons was more hyperpolarized in FXS slices than in WT slices (Figure 8; n = 6 cells from 4 rats of each genotype, univariate analysis of variance, genotype as between-subjects factor, significant effect of genotype (F(1,10) = 5.0, p = 0.05)).

Next, we found that TGOT administration affected FXS and WT neurons differently (Figure 8; general linear model with genotype as between-subjects-factor and drug treatment as repeated measure within subjects, significant interaction between genotype and drug treatment (F(1,10) = 5.4, p = 0.04)). Specifically, TGOT application significantly depolarized CA2 pyramidal cells in slices from WT rats (significant difference in post-hoc pairwise comparison for WT rats for baseline vs. TGOT (ΔV_M_: 3.8 ± 0.83 mV; t(5) = −4.6, p = 0.006)), as expected (Tirko et al. 2018). In contrast, CA2 pyramidal neurons in slices from FXS rats did not significantly depolarize in response to TGOT application (no significant difference for FXS rats for baseline vs. TGOT (ΔV_M_: 1.2 ± 0.79 mV; t(5) = −1.472, p = 0.201)). These results suggest that deficient responses to oxytocin in CA2 neurons may hinder activation of CA2 neurons by social stimuli and thereby contribute to deficits in social remapping observed in CA2 place cells in FXS rats.

## Discussion

A large number of studies have identified abnormalities in social behaviors in mouse and rat models of FXS (Hamilton et al., 2014; Liu & Smith, 2009; McNaughton et al., 2008; Mineur et al., 2006; Oddi et al., 2014; Saxena et al., 2018; Spencer et al., 2005; Tian et al., 2017; Wong et al., 2020). In this study, we identified a neurophysiological difference in responses to social stimuli in CA2 place cells of FXS rats. We have previously shown that a significant proportion of CA2 place cells in WT rats changed their firing fields when a familiar rat in a soiled cage was presented (Alexander et al., 2016; Robson et al., 2025) or when a soiled home-cage was presented without a rat (Robson et al., 2025). In contrast, the present study revealed that CA2 place cells in FXS rats did not change their firing fields when social stimuli were presented. This lack of social remapping in CA2 of FXS rats was not explained by reduced exploration of social stimuli or a lack of olfactory discrimination in FXS rats. Furthermore, we found no differences in firing rates, spatial information coding, and basic place field properties in CA2 place cells in FXS rats, suggesting that impairments in CA2 place cell activity in FXS rats were largely specific to responses to social stimuli.

To our knowledge, this study is the first to record CA2 place cells in a rodent model of FXS. We found no abnormalities in baseline firing rates or spatial coding in CA2 neurons in FXS rats. However, our experiments exclusively involved familiar environments and stimuli. Environmental novelty has been shown to differentially affect CA1 place cells in FXS rats (Asiminas et al., 2022). Specifically, CA1 place cells in FXS rats do not undergo normal experience-dependent changes in spatial selectivity or firing rates (Asiminas et al., 2022). Therefore, it is possible that CA2 place cell properties would differ between WT and FXS rats across experience in a novel environment. Future experiments examining CA2 place cell firing patterns in FXS rats exploring novel environments will be important to determine if CA2 place cell firing properties are differentially affected by experience in WT and FXS rats.

Multiple studies have shown that CA2 place cells alter their firing fields during social interactions (Alexander et al., 2016; Oliva et al., 2020; Robson et al., 2025), but the mechanism underlying this social remapping has not yet been identified. Therefore, it is possible that impairments in CA2 place cell remapping in FXS rats may be inherited from upstream regions or may be due to impaired local computations within CA2. CA2 is unique as a hippocampal subregion in that it contains an abundance of neuropeptide receptors, including oxytocin. In mice, blocking or ablating oxytocin receptors in CA2 impairs social memory (Lin et al., 2018; Raam et al., 2017; Tsai et al., 2022), and CA2 neurons are depolarized by activation of oxytocin receptors (Tirko et al., 2018). It is therefore possible that the release of oxytocin in CA2 during social interactions promotes changes in CA2 place cell firing patterns. If so, a dysregulation of oxytocin release, reduced expression of oxytocin receptors, or impaired oxytocin signaling in CA2 of FXS rats could contribute to the lack of remapping to social stimuli observed in FXS rats in this study. In FXS rats in the present study, we identified reductions in oxytocin receptor expression levels and impaired depolarizing responses to oxytocin in CA2 neurons. These oxytocinergic impairments may contribute to deficient place cell coding of social stimuli in the following manner. Depolarizing oxytocinergic inputs to CA2 may offset strong inhibitory inputs to CA2 neurons (Chevaleyre and Siegelbaum, 2010), thereby allowing some CA2 neurons to develop a new place field in response to social stimuli. Considering that the overall level of activity in the place cell network is thought to be homeostatically maintained (Kaleb et al., 2021), newly activated place cells could trigger network competition and local inhibition within CA2, causing some previously active cells to lose their place fields. In this way, activation of oxytocin receptors on CA2 place cells could promote remapping to social stimuli in WT rats. In contrast, diminished oxytocin receptor expression and weakened depolarizing responses to oxytocin could hinder CA2 place cell remapping to social stimuli in FXS rats.

Oxytocin release in CA2 has not yet been characterized in FXS models, but reduced oxytocin release may also potentially contribute to these deficits. Oxytocin receptors are overexpressed in several subregions of the hippocampus, including CA2, in a separate rodent model of autism, potentially as a compensatory mechanism for reduced oxytocin release (Bertoni et al., 2021).

Previous work has also shown that the oxytocin-mediated excitatory/inhibitory shift of GABA is impaired in a mouse model of FXS (Tyzio et al., 2014), and that treatments that rescue this impairment prenatally result in improvements in social behavior in adulthood (Eftekhari et al., 2014). Related to this, an important question to be examined in future studies is whether oxytocin administration (e.g., oxytocin uncaging; Ahmed et al., 2023) can rescue CA2 place cell remapping in FXS rats.

CA2 receives direct inputs from other brain regions involved in social processing that may be important for transmitting social information to CA2 and driving social responses in place cells. The direct projection from the lateral entorhinal cortex to CA2 is necessary for social memory (Dang et al., 2022; Lopez-Rojas et al., 2022). This direct projection arises from layer II entorhinal cortical neurons (Dang et al., 2022). To our knowledge, layer II entorhinal cortical neurons have not been examined in a rodent model of FXS, but neurons in other layers of the entorhinal cortex show abnormal physiological properties in FXS mice (Deng et al., 2013; Deng & Klyachko, 2016). Thus, an interesting hypothesis to test in future studies is the extent to which abnormal physiology in the direct projection from the lateral entorhinal cortex to CA2 contributes to impaired CA2 place cell remapping to social stimuli in FXS rats.

The function of CA2 place cell remapping to social stimuli remains unclear. Dorsal CA2, the area from which the place cells in this study were recorded, projects directly to ventral CA1 (Meira et al., 2018; Okuyama et al., 2016), and this pathway is essential for social memory (Meira et al., 2018; Tsai et al., 2022). Exogenous activation of this pathway rescues social memory deficits in a mouse model of autism (Cope et al., 2023) and in an oxytocin receptor knockout mouse (Tsai et al., 2022). Also, the direct projection from ventral CA1 to the prefrontal cortex is important for social memory (Phillips et al., 2019; Sun et al., 2020). Hyperexcitability in this pathway has been shown to contribute to social memory impairments in a mouse model of autism (Phillips et al., 2019). While this pathway has not been specifically examined in FXS models, FXS mice show reduced coherence between the ventral hippocampus and the prefrontal cortex (Bhandari et al., 2024), providing a potential mechanistic link between impaired CA2 place cell remapping in FXS rats and abnormal social behaviors.

However, the possibility remains that CA2 place cell remapping to social stimuli is epiphenomenal. CA2 place cells also remap to the presentation of novel objects (Alexander et al., 2016; Oliva et al., 2020), but optogenetic silencing of CA2 neurons does not impact performance on a novel object recognition task (Oliva et al., 2020). Although inactivation of CA2 impairs social memory (Hitti & Siegelbaum, 2014; Leroy et al., 2017; Oliva et al., 2020; Stevenson & Caldwell, 2014), it is difficult to establish if the change of firing fields in response to social conditions is necessary for social memory or if a more general CA2 signal is sufficient. It is possible that the impaired CA2 remapping to social stimuli in FXS rats may not functionally contribute to abnormal social behaviors. Future studies are necessary to establish whether CA2 place cell remapping has functional consequences for social behaviors and, if so, how impairments in CA2 place cell remapping may contribute to abnormal social behaviors in FXS rats. Establishing these connections may help provide novel treatment targets that specifically address social behavioral impairments in patients with FXS.

## Acknowledgments

This research was supported by the Department of Defense CDMRP award W81XWH1810314 (to L.L.C.), Brain and Behavior Research Foundation (BBRF) Distinguished Investigator Grant 32221 (to L.L.C.), and National Institutes of Health awards R56MH125655 (to D.H.B. and L.L.C), R01MH131317 (to D.H.B. and L.L.C.), and F31MH127933 (to M.M.D.). The authors thank Isabella Lee, Ayomide Akinsooto, and Sirisha Dhavala for technical assistance and Chenguang Zheng for providing MATLAB code for some of the analyses. We also thank Dr. Moses V. Chao and Dr. Robert C. Froemke from New York University School of Medicine for generously providing the OXTR-2 antibody used in this study. The authors acknowledge the Texas Advanced Computing Center (TACC) at The University of Texas at Austin for providing data storage resources that have contributed to the research described within this article. URL: http://www.tacc.utexas.edu

## Author contributions

M.M.D. performed research, analyzed data, and wrote the paper; E.R. designed research, performed research, analyzed data, and wrote the paper; A.M.M. performed research; E.J.F. analyzed data; M.H. performed research and analyzed data; A.J.M. designed research, performed research, and analyzed data; J.B.T. performed research; D.H.B. designed research, performed research, and wrote the paper; L.L.C. designed research, analyzed data, and wrote the paper.

---

**Figure S1.**
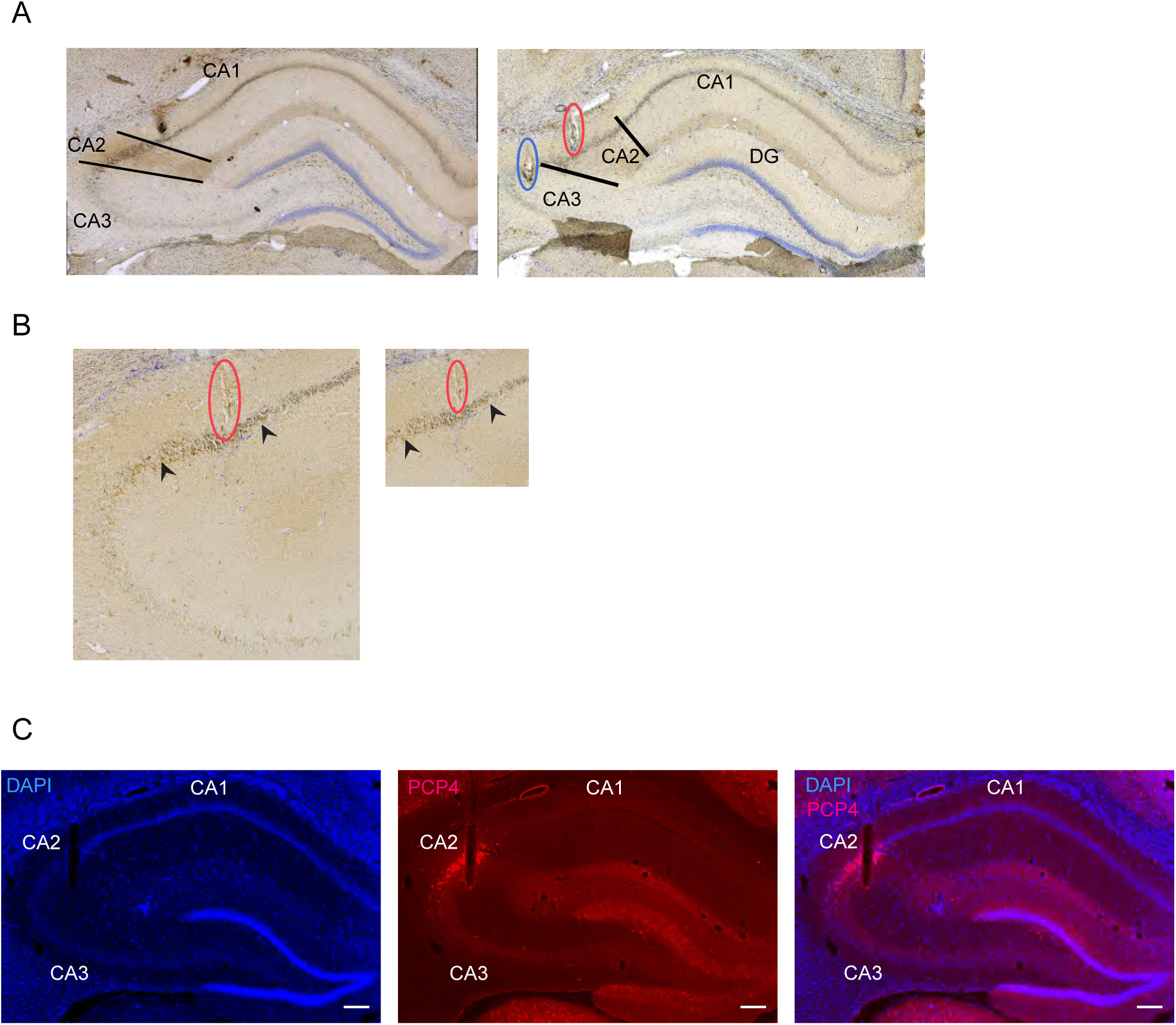
Histological identification of CA2. (A) In four rats, brains were immunostained with the CA2 marker STEP (brown) and cresyl violet (purple). The boundaries of CA2 are denoted by black lines in both images. Subregions CA1, CA2, and CA3 are labeled. Right: Two example tetrode tracks are shown. The tetrode circled in red recorded place cells from CA2, and the tetrode circled in blue recorded place cells from CA3. (B) In rats in which STEP staining did not clearly differentiate the boundaries of CA2, identification of CA2 was aided by its characteristic increase in cell body size. Boundaries of CA2 are indicated by the black arrows. A CA2 tetrode track is circled in red. (C) In eight rats, brains were immunostained with CA2-expressed protein (Purkinje cell protein 4 (PCP4); red) and DAPI (blue). A CA2 tetrode track is shown.

**Figure S2.**
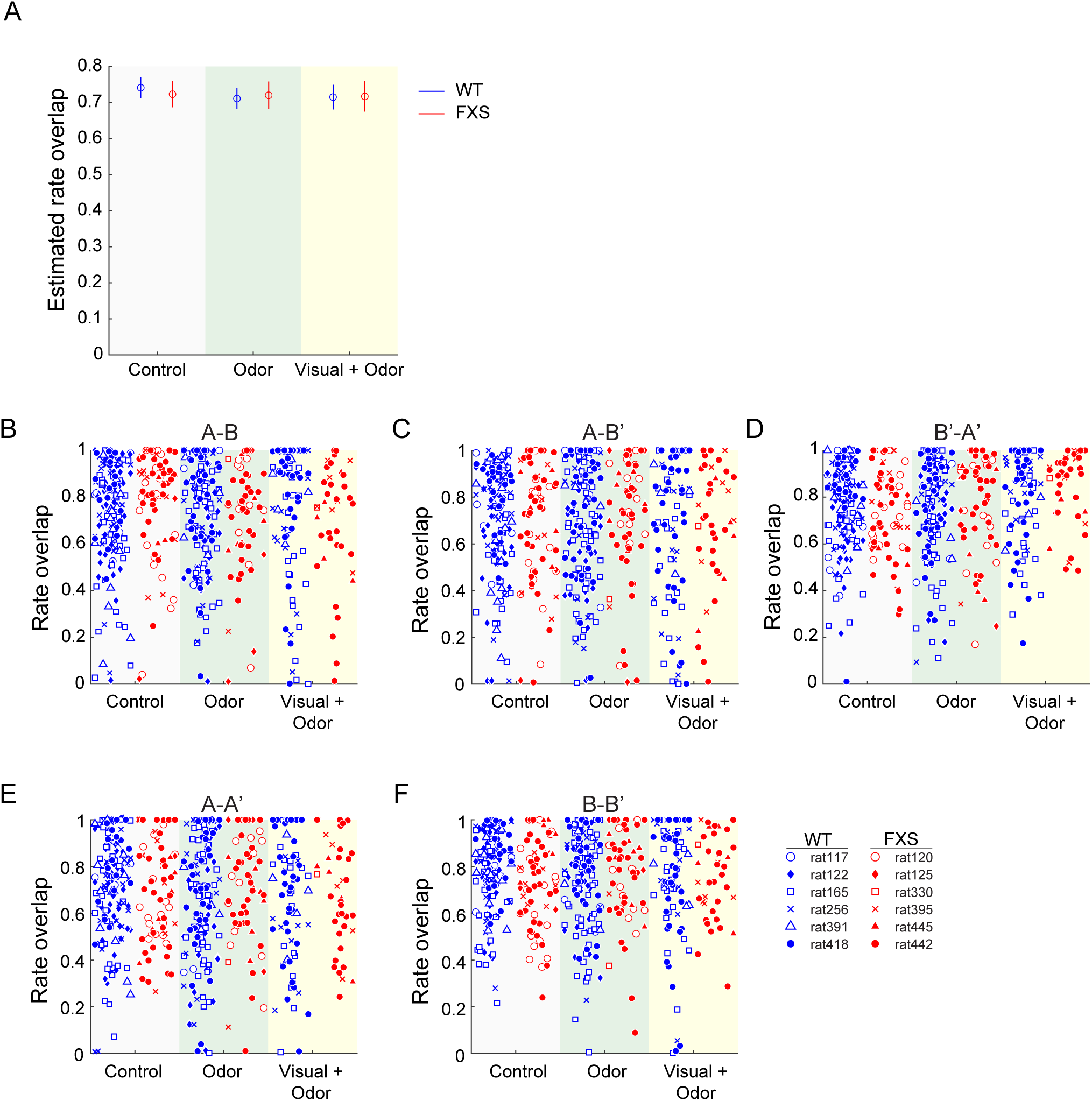
CA2 place cells did not significantly alter their firing rates in response to social stimuli in WT or FXS rats. **(A) The estimated means of rate overlap values from a generalized linear mixed model are shown for each genotype across each condition for all session pairs combined. Dots represent estimated mean values, and error bars represent the estimated 95% confidence intervals. Rate overlap values did not differ across genotypes or conditions. (B-F) Rate overlap values are shown for the entire sample of CA2 place cells for all pairs of sessions across all conditions. Each marker represents a rate overlap value for an individual place cell. Different symbols are used for CA2 place cells recorded from different rats. Values from WT and FXS rats are shown with blue and red symbols, respectively.**

